# Aggregation dynamics of terrestrial isopods under extended Vicsek model

**DOI:** 10.64898/2026.10.09.757868

**Authors:** Pablo Vidal-Franco, Marcin Zagórski

**Affiliations:** Doctoral School of Exact and Natural Sciences, Jagiellonian University, Cracow, Poland; Institute of Theoretical Physics and Mark Kac Center for Complex Systems Research, Jagiellonian University, Cracow, Poland

## Abstract

Terrestrial isopods display a rich behavioral repertoire, including fast escape responses and aggregation dynamics. Their response to environmental stimuli such as artificial light, humidity, vibration, heat and predation stress has been documented, but research highlights a variability in these responses, depending on the studied species and the different ecological niches they inhabit. The Vicsek model is widely used in the study of collective behavior and has been adapted multiple times to model different biological phenomena. In this work, we exposed terrestrial isopod *Porcellionides pruinosus* to different combinations of vibration and light stimuli within a circular enclosure and used the resulting experimental observations to extend the Vicsek model to account for aggregation dynamics and speed variability. To this end, we modeled aggregation dynamics using local group size and edge proximity to modulate aggregation probability. Our experimental results suggest that the combined effect of light and vibration increases the speed of isopods more than light or vibration alone. Additionally, the speed relaxation also occurs later when exposed to the combined stimuli. Interestingly, the classic Vicsek model correctly predicted the motion of isopods only within a few seconds under stimuli presence, indicating the isopods move in a collective fashion at short time scales. In contrast, the results of simulating swarms with the extended Vicsek model were consistent with the mean square displacement of the experimentally tracked isopods over longer times and also, depending on the stimuli, for the short times. This extension of the Vicsek model provides a framework that can be used for multi-modal stimuli testing of terrestrial collective and aggregation invertebrate behavior, thus providing a platform to understand better the variability of the responses of these valuable members of the ecosystem.

**Author summary:** We examined the behavior of terrestrial isopods, focusing on their speed and group dynamics. Terrestrial isopod reactions to environmental factors like light, humidity, vibration, heat, and predator stress are known, but they vary between species and niches. We extended a model that has been widely used for studying collective behavior, to include aggregation dynamics based on terrestrial isopods’ behavior under different combinations of vibration and light stimuli. We found that the combination of light and vibration increases isopod speed more than either alone. We also found that speed relaxation takes longer when both stimuli are present. Finally, isopods tended to move fast in collective fashion for short times but formed still aggregates at longer times after stimuli. Together, this indicates that the levels of stress response for isopods can accumulate and last longer when different stimuli are combined.

## Introduction

Terrestrial isopods have developed a wide range of behavioral adaptations to land life such as stridulation, fast escape responses, conglobation, tonic immobility and aggregation (1,2). Responses to mechanical stimuli have been considered in recent years. A study in *Armadillidium officinalis* tested both stridulating vibration patterns and substrate-borne vibrations and found that the reaction to substrate-borne vibrations was significantly higher (3). There is variability in the way different terrestrial isopods species respond. Wild *Armadillidium officinalis* and *Armadillidium granulatum* subjects showed longer tonic immobility than captive ones when being exposed to predation cues. On the other hand, the response length of wild *Armadillidium vulgare* did not differ from captive ones, suggesting that predation influences these behaviors (4). This variability also appears in other species: when exposed to a combination of heat and vibration stress, *Porcellio scaber* and *Porcellionides pruinosus* differ in the dynamic and size of the aggregates they form (5). In *Porcellio laevis*, single isopods exposed to a car alarm actively avoided the speaker and increased their speed (6).

Earlier studies researching terrestrial isopods’ response to light highlighted their negative phototaxis (7). Recently, their response to this type of stimulus has also been quantified. When exposed to fluorescent and ultraviolet light, *A. vulgare* and *Armadillidium nasatum* displayed increased aggregation (8). It has been shown that the spectral composition of artificial light at night (ALAN) affects the activity patterns of *Tylos spinulosus*, indicating that light pollution can have an impact on these species (9). Similarly, ALAN increased tonic immobility length and shelter-seeking behavior in groups of *P. pruinosus* (10). Despite their aversiveness to light exposure, a mixture of wild *A. vulgare, P. scaber* and *Oniscus asellas* also showed their ability to learn that a well-lit space is associated with (beneficial) nesting material (11). A field study with *P. scaber* and *Armadillidium versicolor* confirmed that these species remained outside artificial shelters mainly during the night (12), which could be related to relative humidity, favorable light conditions, or a combination of both.

The aggregation and sheltering dynamics of terrestrial isopods have also been studied. A population of cave-adapted *Asellus aquaticus* has been compared against a population of surface-adapted conspecifics. It was found that surface-adapted individuals sheltered more frequently whereas aggregation differed between populations of either adaptation (13). Another form of sheltering which results in no movement is conglobation. It has been found that the time it takes to unroll after being exposed to predation stimuli is significantly larger in males (14). It has been theorized that aggregation may have a social component, on account of *A. vulgare* individuals choosing to aggregate under a shelter where other isopods had already aggregated (15).

On the theoretical and computational side, the collective behavior in animals is often investigated with agent-based flocking models that retain information about individuals (16). One of such models, based on finding consensus direction of movement for group of agents within predefined interaction region is the Vicsek model (17). The simplicity of the Vicsek model makes it fundamental for the study of collective motion (18). Earlier modifications on the Vicsek model explored particle synchronization via partial communication and have found that a low communication cost is sufficient to produce cluster aggregation (19); additionally, different tiers of leadership potential have been discussed in the Vicsek model, producing different distribution patterns (20). In the context of cells motility, the modification of the Vicsek including short-range interactions successfully reproduced swarm-like behavior of keratocytes placed on a microfabricated arenas (21), and experimentally the local alignment and force transmission were identified as factors in collective cell migration (22). Other extensions included concept of self-propelled interacting particles as tissue models, indicating cellular motility and adhesion, as factors affecting whether cells in the tissue move or remain in an arrested state (23). The Vicsek model has been widely developed in recent years, including improvements to make agents reach motion consensus faster (24,25). Extensions to the model to reproduce biological behavior more faithfully have also been proposed: e.g. delayed signal propagation that increases swarm robustness to noise (26) and competing agents that produce parallel and anti-parallel swarming (27). The Vicsek model has also been used as a base to model collectives’ trade-off between consensus and responsiveness in complex environments (28). In the context of insect swarms, a harmonically confined version of the model has been found to have two distinct chaotic regimes (single- and multi-cluster) which are separated by a scale-invariant phase behavior (29,30).

Here, we collect experimental data by tracking terrestrial isopods subject to different combinations of vibrations and light stimuli in controlled conditions. We quantify this data in terms of speed and spatial occupancy, and verify against predictions of the Vicsek model restricted to circular arena. Although the model correctly reproduces short-time dynamics of isopods under most of the conditions, it already fails after a few seconds to explain the spatial and temporal behavior of the isopods at longer times. As we show, the observed behavioral responses of terrestrial isopods to stimuli indicate the formation of non-moving aggregates of isopods. These aggregates almost uniquely form near the boundary of circular arena suggesting that isopods are finding shelter in these regions. Combining these observations, we extended the Vicsek model to include aggregation as stochastic process in which simulated isopods have tendency to form aggregates based on proximity to other isopods as well as the distance to the edge of the arena. The results of the extended model are in good agreement with both the speed and spatial experimental data, providing better understanding of aggregation dynamics of terrestrial isopods in response to light and a set of configurable vibration stimuli.

## Materials and Methods

### Animals

Swarms of 25 *P. pruinosus* individuals bought in a local shop specializing in arthropods were used for this experiment. The sampled populations only included male/female adults and juveniles (*mancae* were omitted), taken at random from the enclosure.

### Experimental setup

The Quadrant Actuator Machine (QAM), a machine that was designed and assembled for this experiment, was used to perform experiments on the model organism to gather data to extend the Vicsek model (31), Fig. 1A. S1 Table shows a summary of the experimental setup. When the lamp was off, an average of 57 lux were measured. When the lamp was on, the number of lux increased to 598. Both measurements were done by a Benetech GM1010 luxometer by measuring 10 random points of the working surface. A Benetech GM320 infrarred thermometer was used to ensure that the temperature of the experimental setup remained within 21 °C − 22 °C. The distance from the camera to the Petri dish was set at 30 cm. The tested swarm size was 25 individuals. Swarms were tested 10 times under every condition. During each repetition, two minutes were reserved for acclimation to the new environment before the quadrant actuators and/or the light were turned on. The actuators tap the Petri dish at 4 hits/s. The total duration of each repetition was 3 minutes. Before each repetition, the Petri dish was wiped with 70% EtOH, then wiped with paper towel and left to air dry for 5 minutes. After each experimental group was tested, the Petri dish was replaced with a new sterile one.

**Fig 1.**
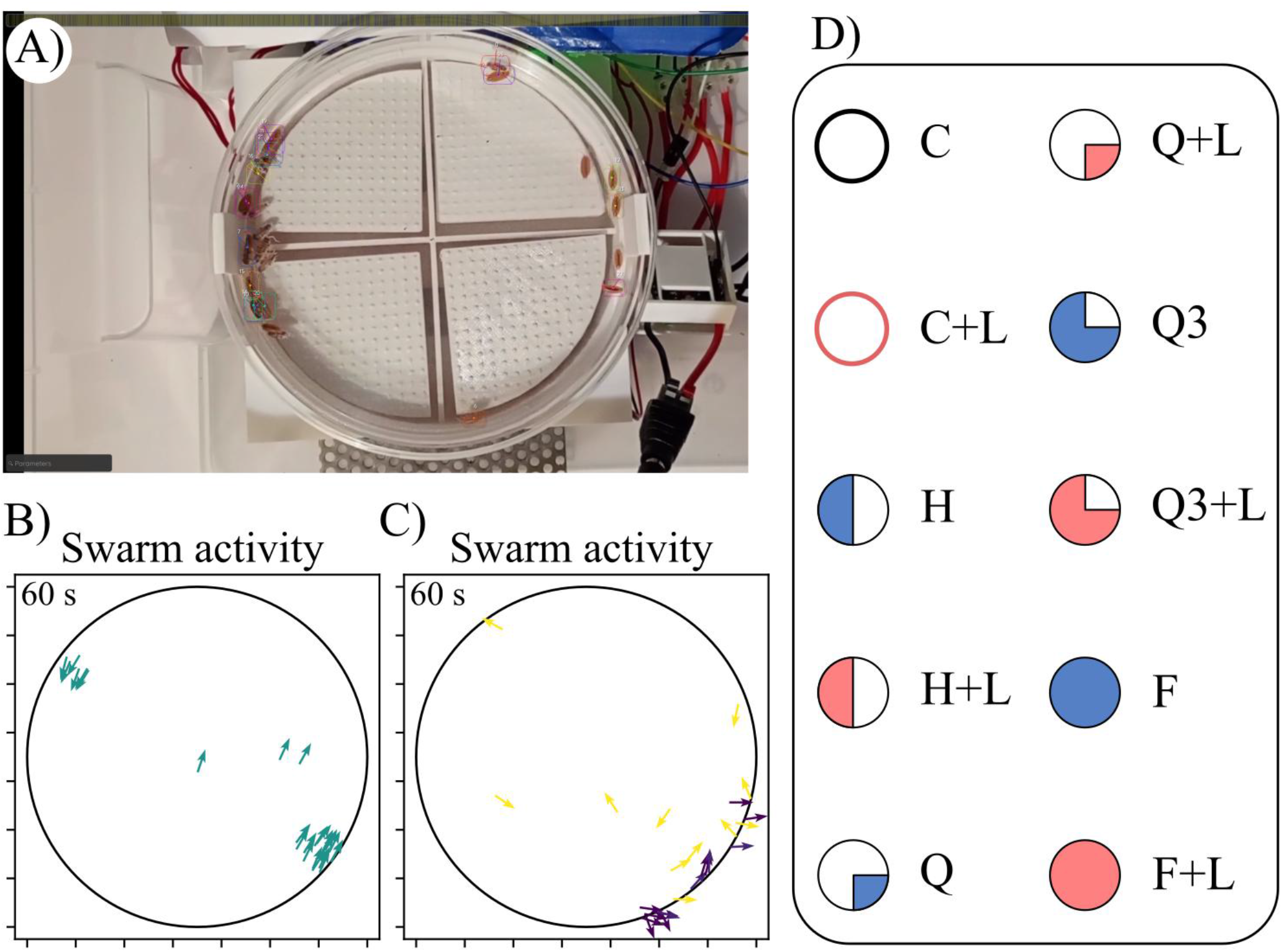
Overview of the experimental setup. **A)** The Quadrant Actuator Machine (QAM) with Petri dish placed directly above the four quadrant actuators. The actuators can be turned on independently. Here, n = 25 terrestrial isopods are present and are tracked by the software TRex (small dots on each of the isopods). **B)** Snapshot of the radially restricted Vicsek model (null model) simulation. Simulated isopods (agents) are represented as arrows, brighter colors correspond to higher speeds. **C)** Snapshot of the radially restricted Vicsek model with aggregation dynamics (extended model) simulation. **D)** Legend with tested conditions: C = control; H = half of the plate actuates; Q = only one quadrant is actuating; Q3 = three quadrants are actuating; F = all four quadrants are actuating. Any condition followed by “+L” indicates that light stimulus is also active. The actuating quadrants are indicated by colored quadrants in the pictograms.

### Motion analysis

The videos were trimmed to the third minute and cropped to fit only the experimental arena. Tracking was done using TRex software (32) with a custom Yolo model, based on the existing yolo11n.pt, trained on annotated frames extracted from the videos (33). A mask was set around the circular arena to ignore tracking out of the region of interest, Fig. 1A.

The data was exported from TRex and analyzed in Python using libraries NumPy, Pandas and statsmodel (34–36). As isopods aggregate or leave an aggregate, loss of tracking momentarily occurs, resulting in NaN values. To mitigate this, a rule was enforced to make sure that every TRex.csv contained at least 75% of non-NaN values. To understand how the speed of the different groups differ at the time that the stimuli are presented, speed snapshots of individual isopods were taken across conditions and compared with a pairwise Welch t-test. These snapshots were taken at *t* = 2 s where *t* = 0 s is the start of the stimuli. Additionally, an exponential curve was fit to the average speed of isopods estimated based on speed snapshots every 0.5 s during the 3^rd^ minute (from *t* = 0 s to *t* = 60 s) in every condition to observe the relaxation of the speed. The log(τ) values of individual isopods in all repetitions per condition were tested with a pairwise independent t-test. Kernel Density Estimations (KDE) of the (*x, y*) coordinates were plotted as motion heatmaps. All plots were produced using matplotlib (37).

### Null model: Radially restricted Vicsek model

The null model is a discretized Stochastic Differential Equation based on the classic Vicsek model (17). The position ***x***_*i*_(*t*) of agent *i* at time *t* is moved by a constant speed v_0_ in the heading angle θ_*i*_(*t*):

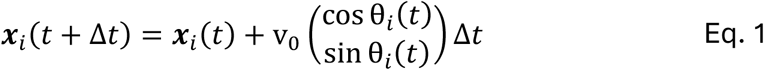

The new direction of the agents is updated with the average heading of their local neighbors based on an interaction radius *R*_inter_, with added uniform noise:

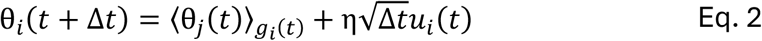

where 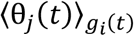 is the average heading angle of agent *i* and its neighbors within radius *R*_inter_, formally 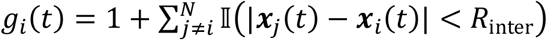 with I(condition) as indicator function, η ∈ [0,2π] as magnitude of noise, and *u*_*i*_(*t*) drawn uniformly from (−1, 1).

Additionally positions of all agents are restricted to a disk (circular arena) of radius *R*, with point (0,0) corresponding to the center of the arena, and *r*_*i*_(*t*) = ||***x***_*i*_(*t*)|| ≤ *R* as radial Euclidean distance of agent *i* from the center. When Eq. 1 is integrated, the direction of agent *i* upon hitting the boundary is handled by first correcting its position such that ***x***_*i*_ ← *R****n***_*i*_ where ***n***_*i*_(*t*) = ***x***_*i*_(*t*) / *r*_*i*_(*t*) is the unit radial direction from the center; and then updating its velocity ***v***_*i*_ ← ***v***_*i*_ − 2(***v***_*i*_ ⋅ ***n***_*i*_)***n***_*i*_ to produce a specular reflection against the boundary of the arena. The new heading angle θ_*i*_ is updated accordingly.

The representative snapshot of collective motion for radially restricted Vicsek model, Fig. 1B. The initial agents’ positions were uniformly distributed over the disk, and their initial heading angles had a uniformly sampled direction from 0 to 2π.

### Extended model: Radially restricted Vicsek model with aggregation dynamics

In experimental observations, the isopods were more likely to stay longer in larger aggregates close to the edge of the circular arena, rather than in smaller aggregates located further from the arena boundary. In order to account for this behavior, we extend the Vicsek model:

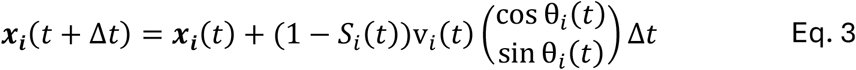

by stochastic state variable *S*_*i*_(*t*) = 1 for agents (isopods) *in the aggregate*, and 0 for agents *outside of the aggregate*. We assume that the *S*_*i*_(*t*) are given by Time-Inhomogeneous Markov Chain (38), so that the current state of the agent depends only on its previous state. The transition probabilities are then following:

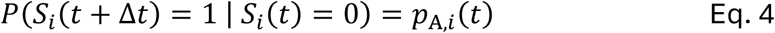

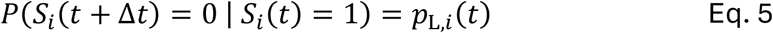

where *p*_A,*i*_(*t*) is probability of joining an aggregate and *p*_L,*i*_(*t*) is the probability of leaving an aggregate for agent *i*. The two remaining transition probabilities are given by normalization, so 1 − *p*_A,*i*_(*t*) for remaining in the aggregate (agent remains still), and 1 − *p*_L,*i*_(*t*) for remaining outside of the aggregate (agent remains in movement). Here, we define:

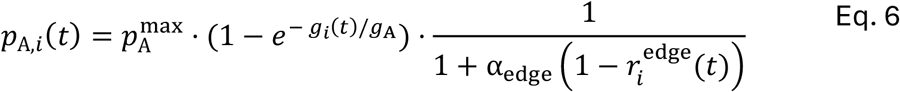

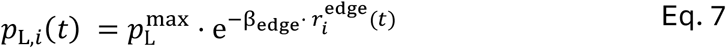

where 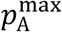 and 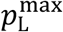 are the per-step maximum probability of aggregating and leaving an aggregate, g_A_ is threshold group size, 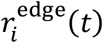 is the scaled normalized distance of agent *i* from the center, α_edge_ is parameter that controls the preference of agents to form aggregates near the edge, and β_edge_ controls the preference of agents to leave aggregates far from the edge. The scaled normalized distance is defined as 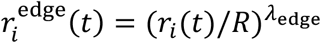 where *r*_i_ (t) is the radial distance of agent *i* from the center of the arena of radius *R*, and exponent λ_edge_ controls the non-linearity, by default λ_edge_ = 3. In practice, we use effective real-valued parameter λ_A_ = 1/*g*_A_ ≈ 1/4, In general, λ_A_ can take values from 0 (no effect from group size) to 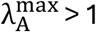 (strong preference for the group size). Furthermore, the 1/β_edge_ can be interpreted as characteristic scaled distance from the edge at which the effect of leaving the aggregate diminishes. Here we set β_edge_ = 0.92, thus 1/β_edge_ > 1, so the probability of leaving the aggregate strongly depends on 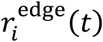. The time dependence of probabilities *p*_A,*i*_(*t*) and *p*_L,*i*_(*t*) enters only through positions of agents and resulting neighborhood sizes, the model parameters are constant in time. The parameters were selected based on the observations made during the analysis of the videos. As isopods seemed to be more likely to aggregate than roam freely, the maximum per-step probabilities to aggregate 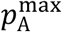 and dissociate 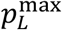 were chosen as the main values to encourage aggregation and discourage dissociation (0.75 and 0.25, respectively). Additionally, aggregation typically happened closer to boundaries than at the center of the arena, thus we chose a large value for α_edge_ = 20, as lower values produced accumulations near the edge instead of at the edge. Summary of all model parameters is in S2 Table, and the effect of separate terms are plotted in S1 Fig.

The representative snapshot of collective motion for radially restricted Vicsek model with aggregation dynamics, Fig. 1C. The initial agents’ positions were uniformly distributed over the disk, and their initial heading angles had a uniformly sampled direction from 0 to 2π.

### Methods to compare experimental data and simulations

10 simulations using swarms of 25 agents were done using the Vicsek null model and the Vicsek model with aggregation dynamics restricted to circular arena. Each simulation consisted of 3600 single updates of Eq. 1 with *Δt* = 0.033 s and 25 agents. The custom code was written in Python and is available at GitHub. We compared which model approximates the experimental data better by measuring the Time-Averaged Mean Square Distance (TMSD). Because loss of tracking can happen during isopod aggregation/disassociation, the TMSD was computed for each tracklet *j* defined as a finite series of consecutive agent positions 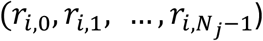 of agent *i*:

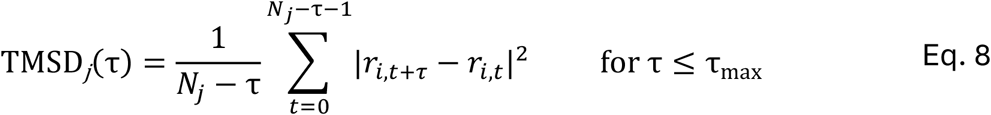

where τ is the time lag between compared positions, *N*_*j*_ is length of the tracklet *j*, and τ_max_ is set to 360 frames, i.e. 12 seconds at 30 FPS. The tracklets are defined such that any two tracklets do not overlap. Tracklet creation was done such that every NaN in the TRex .csv export produced the beginning of a new trajectory (i.e. a tracklet). We considered only tracklets of minimal length *N*_*j*_ ≥ 10. As tracklet length varies, the pooled TMSD was computed as a weighted average of the per-tracklet TMSD:

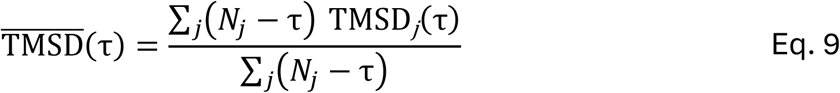

Another way to compare experimental data and models was to visualize the spatial occupancy of isopods/agents. To do so, a 2D Kernel Density Estimate (KDE) map was fit on the 14 cm by 14 cm arena (binned in a 60 by 60 lattice) using the positional data of the 25 isopods only in the minute of exposure to stimuli with data of every repetition of a given experimental setup pooled together. The bandwidth parameter governing the standard deviation of the Gaussian kernel was set to 0.4 cm, approximately a quarter of an isopod length. To this end, the KernelDensity class of library sklearn was used with kernel parameter set to “gaussian” and bandwidth parameter set to 0.4 cm (39). In all cases, speeds over 8 cm/s were regarded as motion tracking artifacts and discarded from the analysis, see Table S3 for statistics of the discarded speeds per condition.

## Results

### Speed of isopods is affected by vibration and weakly by light stimuli

First, we wanted to test whether switching on light without vibrations results in increased speed of isopods. We collected isopod speeds shortly after the light activation and found that there is no significant difference (pairwise Welch t-tests, Holm-Bonferroni adjusted p-value (p-holm) ≥ 0.05) between isopods average speeds upon stimuli activation between control condition (C), 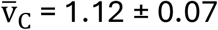 (mean ± SEM), and control with light condition (C+L), 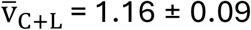 (mean ± SEM), Fig. 2. The distributions of isopod speeds were also only slightly broader for conditions with light activated *σ*_v,C+L_ = 1.19 cm/s than in the condition without light *σ*_v,C_ = 1.03 cm/s. Hence, we concluded that without vibration switching on, light does not affect isopods behavior at short time scales.

**Fig 2.**
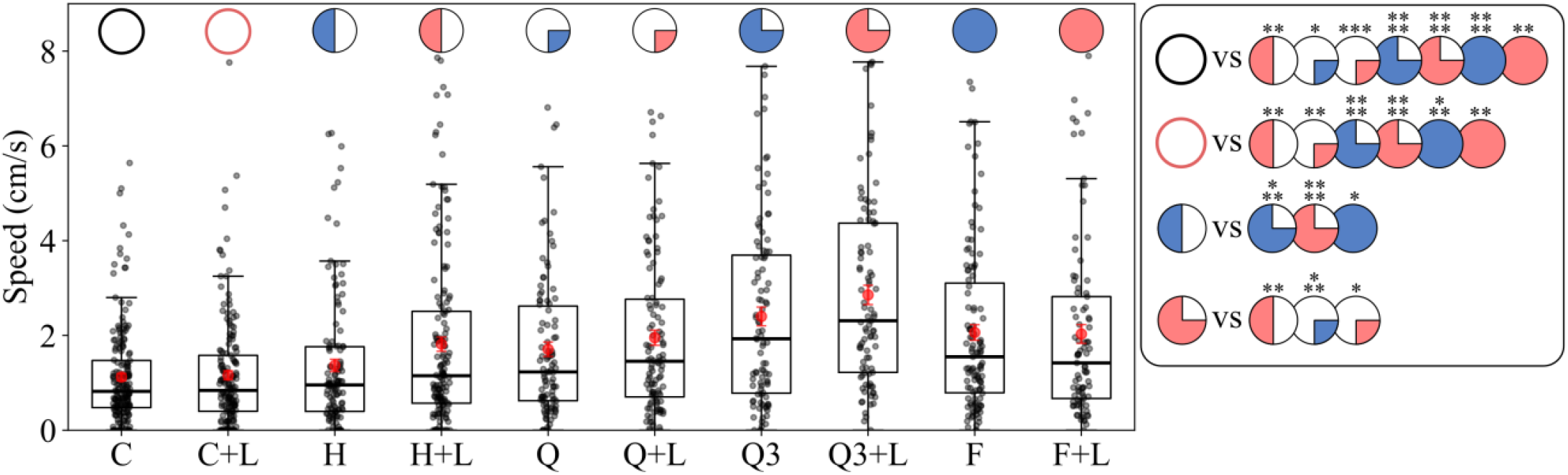
Speed of isopods shortly after the stimuli. Tracked speed of isopods at *t* = 2 s after stimuli activation. Each dot corresponds to the recorded speed of single isopod. Different conditions with sample sizes: C (control, n=188); C+L (control + light, n=161); H (half-vibrating, n=120); H+L (half-vibrating + light, n=153); Q (quadrant-vibrating, n=102); Q+L (quadrant-vibrating + light, n=110); Q3 (three quadrants vibrating, n=96); Q3+L (three quadrants vibrating + light, n=93); F (full-vibrating, n=115); F+L (full-vibrating + light, n=89). The right-most plot shows significance for pairwise Welch independent t-test between different conditions. Significance: **** p < 0.0001; *** p < 0.001; ** p < 0.01; * p < 0.05. Non-significant values (p ≥ 0.05) are not shown. See Fig. 1D and S1 Table for specification of the experimental setup. The mean (red point), median (thick black line), lower/upper quartile (box boundaries), lowest/highest data point within 1.5·IQR above the lower/upper quartile (whiskers) are indicated.

Next, we investigated to what extent the C and C+L conditions differ from conditions with different levels of vibrations and when vibrations were combined with light activation. We found that switching on vibrations significantly affects the average speed of isopods for most of the conditions (all except for half-vibrating (H)). Specifically, as quantified by the pairwise Welch t-tests, the largest difference is against condition with three-quadrants vibrating (Q3), three-quadrants with light stimuli (Q3+L) and the fully vibrating plate (F) (all, p-holm < 0.0001 against C), with 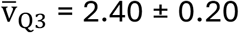 (mean ± SEM), with 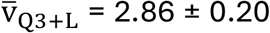 (mean ± SEM) and with 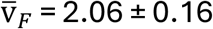 (mean ± SEM), Fig. 2 and S2 Fig. The large differences were also observed, against one-quadrant vibrating with light (Q+L), 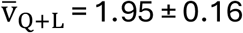 (mean ± SEM) (p-holm < 0.001), and fully vibrating plate with light (F+L), 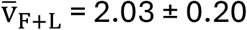 (mean ± SEM) (p-holm < 0.01). In the case of comparing conditions against C+L, the largest difference still occurs against Q3 and Q3+L (p-holm < 0.0001), with a minor drop in significance of F (p-holm < 0.001), and the rest of conditions seeing an attenuation of the significance, Fig. 2 and S2 Fig. Overall, isopods in both control and control with light conditions move significantly slower than in conditions with vibration and vibration combined with light.

Further, we investigated whether isopods under conditions with one or two stimuli present exhibit different speed shortly after stimuli activation. We expected to see this difference for Q3 and Q3+L against other non-control conditions, and indeed the isopods under Q3+L were significantly faster than in H, H+L, and Q, Fig. 2. Interestingly, the isopods in Q3 showed significant difference only against H (excluding controls), and when compared with other conditions with stimuli the difference was insignificant. When investigating speed difference for any other conditions with one (H, Q, F) or two stimuli (H+L, Q+L, F+L) the difference of average speed of isopods, was also not-significant between these conditions, (p-value ≥ 0.05), S2 Fig. We concluded that in most cases already activating vibration makes isopods move faster, and addition of light makes a small difference, with the notable exception of three quadrants vibrating with added light.

### Isopods adapt to different stimuli typically within several seconds

Knowing how isopods are affected by vibration and light stimuli at short time scales, we wanted to understand whether the initial increase of the speed of isopods is maintained over longer times. To do so, we quantified the average speed of isopods every Δ*t* = 0.5 s, for a period of 60 s after initial stimuli (hence 15 frames per bin, 120 bins), Fig. 3. As expected, for C and C+L the speed of isopods was fluctuating around a constant value of 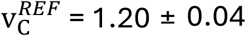 (mean ± SEM) for C, and 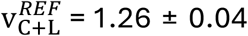 (mean ± SEM), when averaged over the considered period, S3 Fig. However, for all other conditions the average speed of isopods significantly dropped from the initial speed. For the conditions (Q, Q3, F, Q+L, H+L, F+L) the average speed of isopods relaxed approximately to 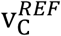 (specifically 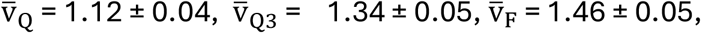,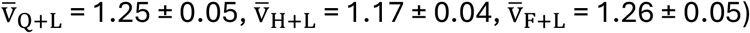. For H the isopods relaxed to a lower speed of about 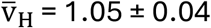 (mean ± SEM) at *t* = 60 s, and for Q3+L, notably, to a higher reference value of 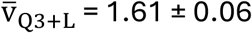 (mean ± SEM) at *t* = 60 s, Fig. 3. Together, this suggests that independent of the combination of stimuli, isopods could relax their speed to the control baseline within 60 s from the initial stimuli, potentially relaxing even faster under different conditions.

**Fig 3.**
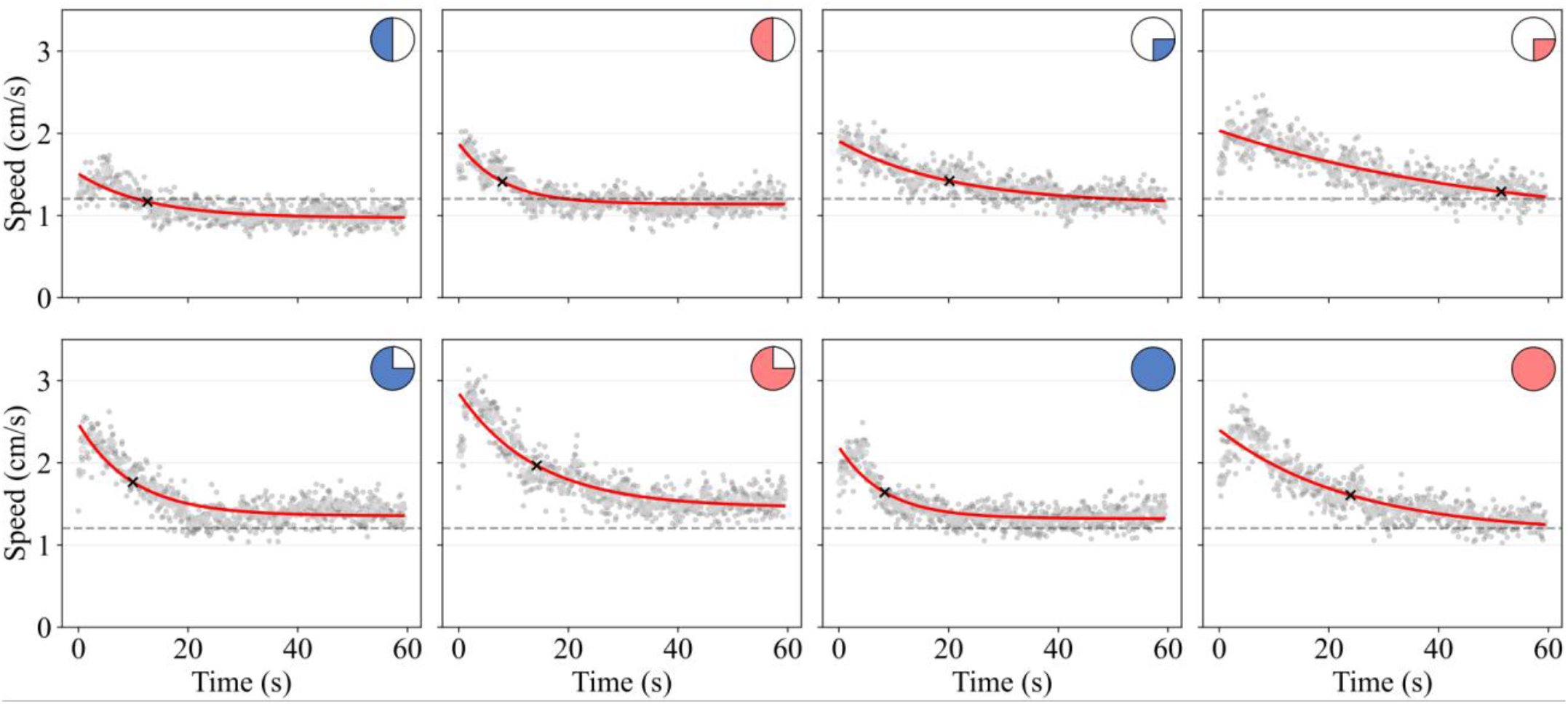
Relaxation time of speed of isopods for different stimuli. The mean speed of isopods (gray points) over 60 s time window, sampled in 0.5 s bins after stimuli activation is indicated. The control mean is displayed as a dashed line. An exponential curve (red) of the form v(*t*) = v_∞_ + (v_0_ − v_∞_)*e*^−*t*/τ^ was fit separately for each condition, where v_0_ was set to the mean speed of isopods in the first bin (*t* from 0 to 0.5 s), and v_∞_ and τ were estimated from the fit. Relaxation times for every tested condition (± SE) are marked with a cross: τ_H_ = 12.6 ± 1.6 s; τ_H+L_ = 7.9 ± 0.8 s; τ_Q_ = 20.1 ± 2.3 s; τ_Q+L_ = 51.4 ± 15.4 s; τ_Q3_ = 9.9 ± 1.0 s; τ_Q3+L_ = 14.2 ± 1.3 s; τ_F_ = 8.3 ± 0.8 s; τ_F+L_ = 23.9 ± 3.2 s. The pictograms indicating different stimuli are as in Fig. 1D.

To test this directly, we fitted exponentially decaying curve with characteristic relaxation time τ to the collected average speeds of isopods within 60 s after stimuli. For most of the conditions (H, H+L, Q3, Q3+L, F) the relaxation time was between 7 s and 15 s, see Fig. 3 for exact estimates. For Q and F+L conditions, the relaxation of isopod speed took slightly longer, τ_Q_ = 20.15 ± 2.33 s, τ_F+L_ = 23.95 ± 3.20 s (mean ± SE). However, for Q+L the fit of exponential function returned τ_Q+L_= 51.42 ± 15.40 s indicating that for this condition the system might require longer time than 60 s to fully relax. When comparing each single-stimulus with its combined-stimuli counterpart (e.g. Q vs Q+L, Q3 vs Q3+L), the relaxation time of isopods speed is always greater for the part with the light activated, except for H vs H+L. Nevertheless, from these pairs, only F vs F+L is statistically significant (p-value < 0.05), see S4 Fig. Altogether, the isopods can relax their speed within several seconds after exposition to the combinations of vibration and light stimuli under most conditions.

### Isopods aggregate more at later times after stimuli

To further understand how isopods adapt to different stimuli we compared their speed distributions for the first and the last 5 seconds for different conditions. For both controls (C, C+L) the speed distributions at early and late times were very similar, Fig. 4. In contrast the same distributions for other experimental conditions showed consistently a clear difference: the higher speeds were overrepresented for the first 5 seconds, whereas very low speeds were overrepresented for the last 5 seconds, Fig. 4. This increase in speed during the first seconds after exposure was especially evident for conditions Q, Q+L, Q3, Q3+L and F+L, which are also all conditions were the relaxation time took significantly longer, see Fig 3. Still, many of the isopods remained in the bulk of the distributions (roughly ≤ 1.6 cm/s). Hence, upon exposure to stimuli isopods disperse relatively fast, yet many of them still move slowly or at moderate speeds. This can be observed especially at the beginning of stimuli activation in Q3+L, which triggers several isopods to rapidly burst across the Petri dish, rather than stay around the edges, see S1 Video.

**Fig 4.**
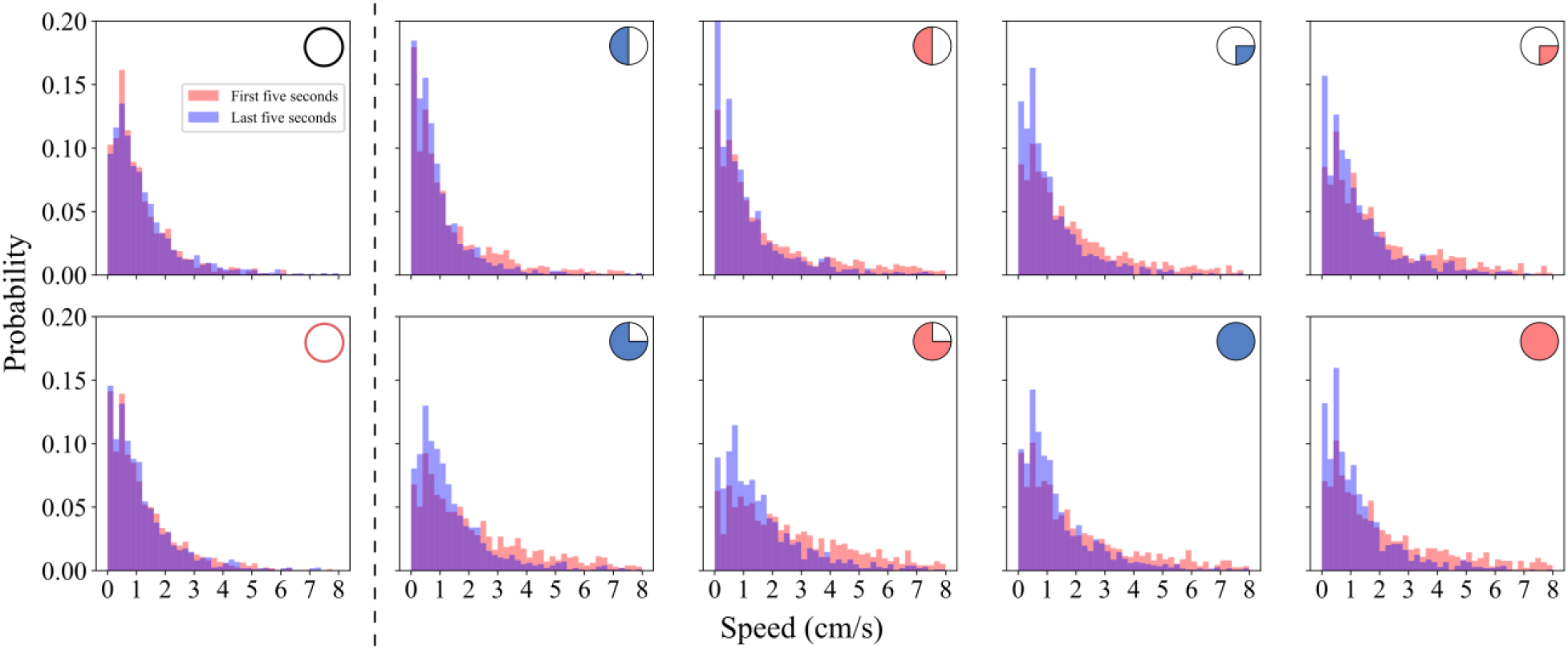
Isopod speed histograms in the experimental conditions after the stimuli activation. The probability distribution of isopod speeds at early times (first 5 s after stimuli activation; red bins) and late times (last 5 s after stimuli activation; blue bins). The control conditions are shown to the left of the vertical dashed line, and the other experimental conditions appear to the right of it. The speed of isopods was sampled from 0.5 s intervals, that is, every 15th frame. The pictograms indicating different stimuli are as in Fig. 1D.

The speed distributions indicated also that at very low-speeds the behavior of isopods is subtle. Specifically, if we focus on the first bin with non-moving isopods (formally with speed < 0.2 cm/s) for the early times, condition H shows a probability of aggregation of about 19%, followed by H+L with 13% and the rest of conditions below 10%. Interestingly, at long times after the stimuli, the probability of finding non-moving isopods is always greater in all conditions. This is particularly visible in H+L (59.80% increase with respect to early time bins), Q (+57.0%), Q+L (+84.22%), Q3+L (+42.15%), and F+L (+86.97%). In the case of H, Q3 and F, the difference is negligible. This increase is also noticeable in the conditions with light relative to their single-stimulus analogues (e.g. H+L vs H, Q+L vs Q, F+L vs F). Finally, we also note a second peak in all conditions between 0 cm/s and 1.0 cm/s range which we speculate could be an indicator of a collective dynamics exhibited by the isopods; in some of the conditions this peak is the main one (C, Q3), and in others it is the highest one present (Q, Q3+L, F, F+L). Taken together, it further supports the concept that many isopods remain still after adapting to stimuli, suggesting that they could form localized aggregates.

To test this hypothesis, we investigated the spatial occupancy heatmaps of the experimental data. The data confirmed that isopods prefer to stay together in well localized hotpots, rather than remain still far from other isopods, Fig. 5. Only for the control, and very weakly for the control with light, the probability of finding isopods in the center of the arena was noticeable. For instance, in S2 Video and S3 Video isopods are crossing the Petri dish more frequently, and the tendency to aggregate directly at the boundaries is less pronounced, with isopods often remaining mobile near the edges instead of fully aggregating. Moreover, in all conditions isopods explored mostly regions close to the boundary of the arena and the probability of finding isopods was highest in a few localized points, indicating preference of isopods to finding shelter near boundary, Fig. 5.

**Fig 5.**
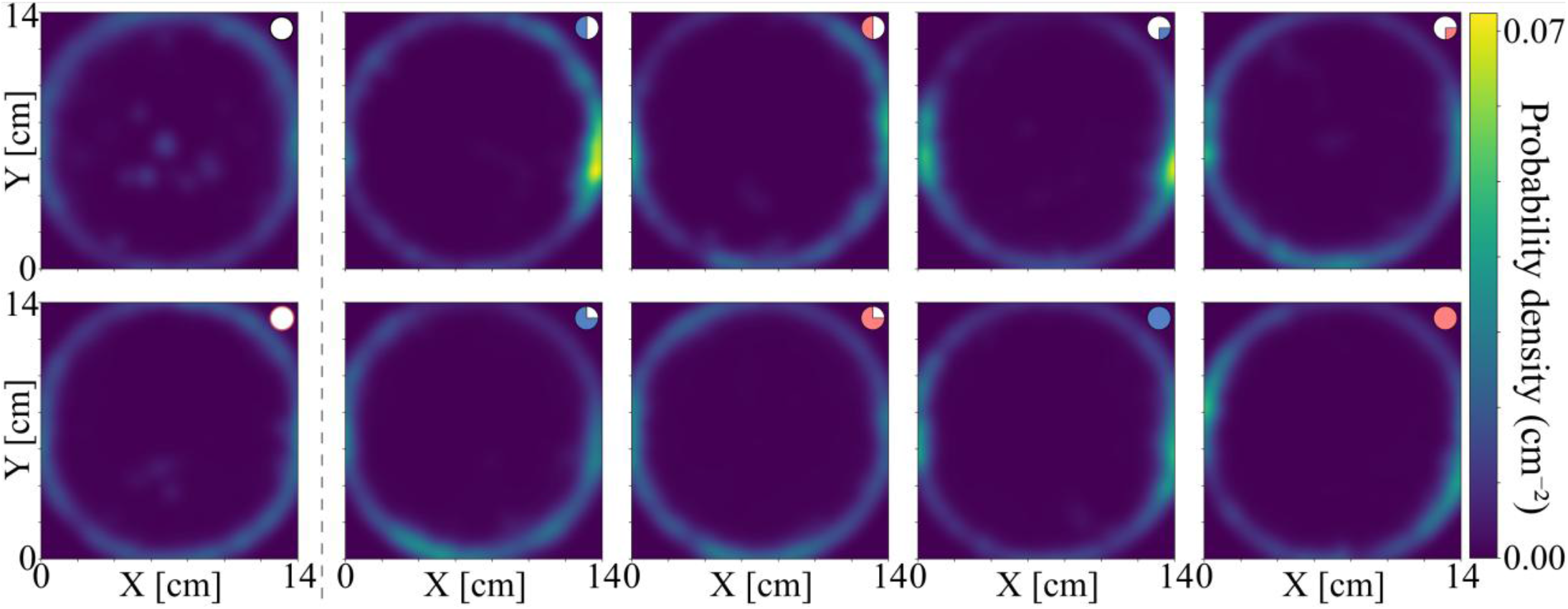
Spatial occupancy of isopods in the experimental data. Spatial occupancy of isopods was quantified as probability density of 2D Kernel Density Estimate map on the 14 cm by 14 cm arena (binned in a 60 by 60 lattice). The positional data of the 25 isopods for each repetition under stimuli presence was pooled together. The control conditions are shown to the left of the vertical dashed line, and the other experimental conditions appear to the right of it.

In order to see whether the observed behavior of isopods can be effectively described by models of collective motion, we first simulated the isopod dynamics with the radially restricted Vicsek model, see Materials and Methods. The simulated isopods (agents) tended to flock together close to the perimeter of the circular arena, however without signs of localization events, Fig. 6. Hence, we extended the Vicsek model by possibility of agents to form spatial localized clusters as a function of number of neighboring agents and as a function of proximity of agents to the edge of the arena, see Materials and Methods, S1 Fig. Specifically, we set the probability of forming an aggregate around a given isopod to be increasing upon more and more agents present within this agent’s interaction radius *R*_*inter*_. As for the preference, to form aggregates near the boundary of the arena, this condition was data-driven, as we wanted to avoid artificial formation of the aggregates in the middle of the arena. In order to control how long isopods spend in the aggregates we also set the probability of leaving the aggregate to be dependent on the proximity to the boundary of the arena, with the lowest value at the edge, and highest in the center of the arena, S1 Fig. As a result, we observed agents to group in specific points along the boundary of the arena. The probability of spatial occupancy of agents in the extended model resembled more closely the experimental data for the control condition. Overall, agents showed tendency to move along the perimeter of circular arena, but most of the time they tended to aggregate in non-moving clusters, Fig. 6. In the remaining experimental conditions after stimuli activation, the circular flocking around the boundaries of the arena was less pronounced and the isopods tended to be localized in one to three aggregation hotspots, as indicated by probability of spatial occupancy, Fig. 5. Taken together, both experimental data and extended model of isopods collective aggregation, suggested the isopods tend to cluster upon stimuli activation more often than in the control condition.

**Fig 6.**
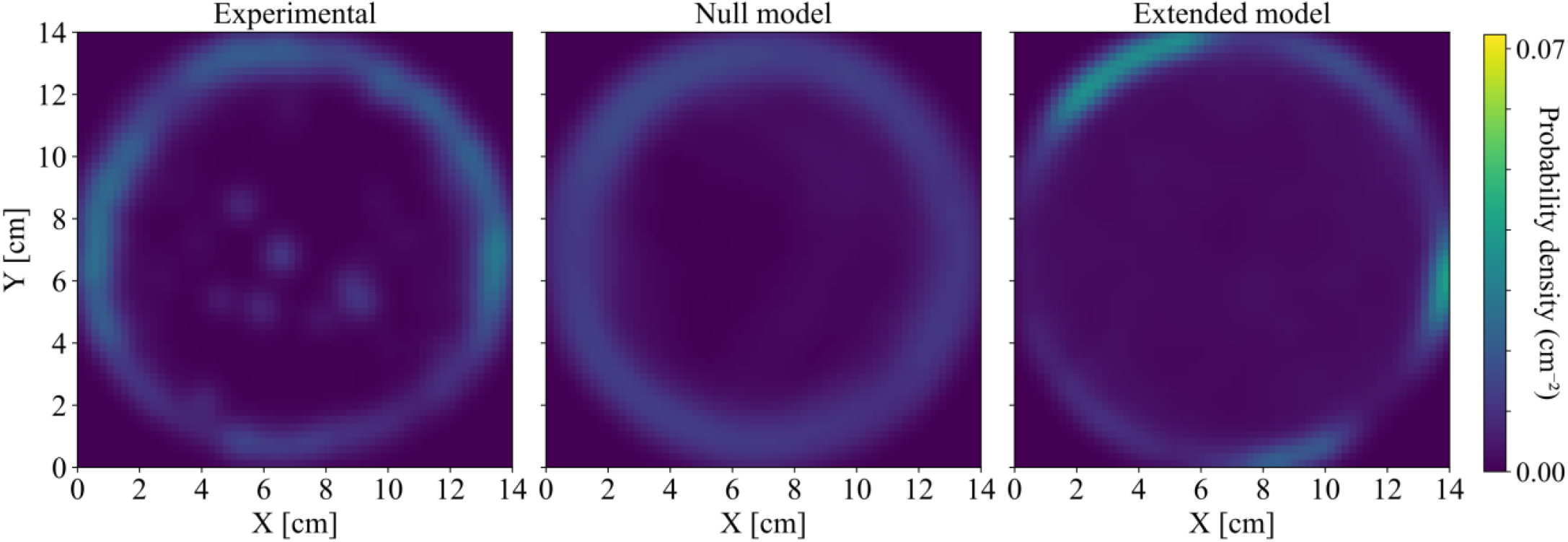
Probability density of spatial occupancy of isopods in the control condition. Left: experimental data KDE using all the repetitions of the isopod positional data aggregated for control condition. Center and right: null model and extended model. Positions of all simulated isopods (agents) were aggregated. See Fig. 5 for the spatial occupancy of the experimental data.

The spatial occupancy data was obtained over long times (1 minute), leaving possibility that for short times (a few seconds), the dynamical behavior of isopods under null and extended model might be inconsistent with the experimental data. To address this point, we calculated time-averaged mean square displacement over tracklets (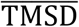, see Materials and Methods) for both models and validated against the experimental data, Fig. 7. The averaged TMSD curves for the null model largely overshoot the experimentally obtained 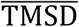 for every condition already after a few seconds, Fig. 7. In contrast, the extended model 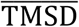 curves stayed within close proximity of the 50% CI bands of the experimental curves, mostly at longer times. In the case of the control, the CI band of both the experimental dataset and the extended model overlapped to a great extent for both short and long times. Despite the extended model outperforming the null model in all experimental setups, we also note that for short times (all except C, C+L, H and Q3+L < 4 s), the null model can approximate terrestrial isopod behavior better. Overall, this indicates that movement of isopods within a few seconds is mostly well described by classic Vicsek model, but at later times aggregation is dominating behavioral response.

**Fig 7.**
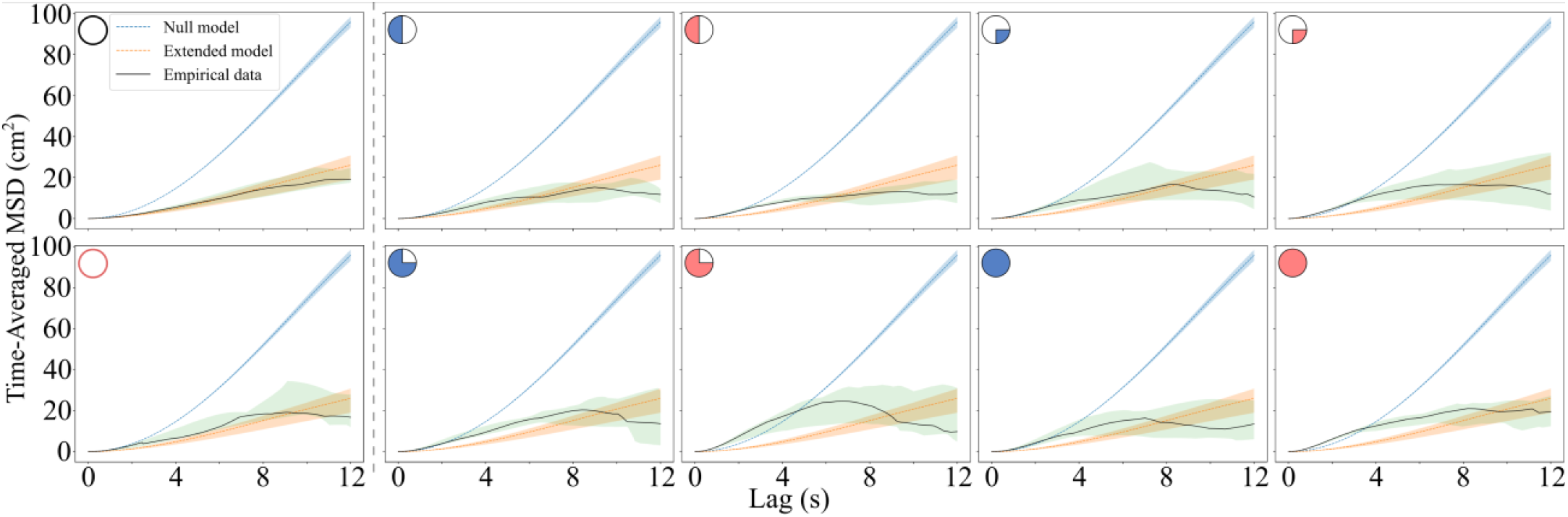
Displacement of isopods in the extended model against the null model and the experimental data for all conditions. The Time-Averaged MSD, 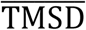, for the null model (blue), extended model (orange), and the experimental data (black, with green confidence bands). The control conditions are shown to the left of the vertical dashed line, and the other experimental conditions appear to the right of it. The bands represent the upper and lower bound quantiles (0.75 and 0.25, respectively). The parameters for null and extended model are as in S2 Table. See Fig. 1D for a description of the pictograms.

Finally, we wanted to understand how these predictions of displacement of isopods over time are affected by modification of parameters present in both null and extended model. First, we note that modifying the speed of agents in both null and extended model by ±50% from the default v_0_ = 1.0 cm/s (the mean for control is 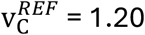), affects monotonically the slopes of respective 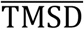, potentially adjusting better with specific experimental conditions for the extended model. For instance, the 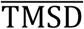 for null model would require half the default v_0_ to match the experimental results, S5 Fig. That is, based on 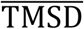 metric, modifying v_0_ in the original Vicsek model would suffice, S6 Fig. However, the spatial occupancy it produces, in a circular boundary, is not representative of terrestrial isopod behavior, S7 Fig. Second, varying the interaction radius of agents from *R*_inter_ = 0.01 cm to 2 cm (the default is 1 cm), resulted in small change of 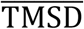 for null model, that could not match the experimental data, S8 Fig. In the null model, a larger *R*_inter_ value has agents average their directions over more neighbors, increasing local alignment and consequently, 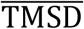. However, the 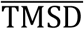 for extended model could be modified in broader range allowing for both results above and below its experimental counterpart, S8 Fig. Particularly, in the extended model, as the interaction radius increases, so does *g*_*i*_(*t*), and thus, the likelihood of aggregation is higher, S1 Fig, which drives 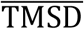 down. Altogether, perturbing speed of agents or their interaction radius does not change the qualitative difference between the two models at longer times, as extended model with aggregation dynamics tends to consistently lower displacements of agents than the null model with only collective motion.

## Discussion

Despite ongoing research on behavioral adaptations of terrestrial isopods to land life, the way terrestrial isopods respond to environmental stimuli, such as subtle mechanical vibrations or light exposure, is poorly understood. We investigated this by assembling a machine with vibrating quadrants and studying spatial and temporal aspects of isopods dynamics in controlled conditions with multi-modal stimuli. By tracking the movement of isopods, we found that isopods exhibit high speed initially, but over time they adapt and are more prone to form localized aggregates than in control condition without stimuli. Further, these aggregates are almost exclusively formed near the boundary of circular arena, potentially indicating need of isopods for finding a shelter. By analyzing the videos of isopods under different stimuli, this tendency to aggregate is present to different extent, with a strong support of sheltering hypothesis in half vibrating condition in which majority of isopods clusters at non-vibrating half, see Fig. 5, S4 Video (H+L). Indeed, the stability of aggregates in terrestrial isopods has been documented to depend on the exposure to stressful stimuli and the number of conspecifics (40). In the control the isopods tend to stop from time to time, forming mostly transient aggregates, S2 Video, and their spatial occupancy is mostly uniformly distributed along the circular boundary of the arena, Fig. 5. These observations are consistent with quantified speed histograms, with many low speed points recorded for conditions with combined stimuli, and relatively few present for control condition, Fig. 4.

In order to understand how much of the isopod motion can be explained by collective dynamics we simulated the swarm of isopods in the circular arena with the Vicsek model (17). The time averaged mean square displacement for isopods simulated in the Vicsek model under the collective movement regime was in good agreement for the first two to four seconds of experimentally obtained TMSD, Fig. 7. Interestingly, this collective motion of isopods for short times was mostly present with combinations of stimuli, and very weakly present for control condition, Fig. 7. This lead us to hypothesize that in the presence of stimuli the isopods feel stressed, and act in more coordinated fashion at short scales, e.g. 3Q+L, S1 Video. After this short response of a few seconds, in all conditions isopods tended to form aggregates, resulting in decreasing the slope of TMSD and flattening of the displacement. We interpret this flattening or even decreasing of the TMSD at longer times as indication that isopods tracked for longer times tend to move relatively little or stop as compared to simulated agents that irrespective of time move in collective fashion. This behavior matches earlier reports in *P. porcellionides*, the subject of this study, where exposure to vibration has been shown to produce more aggregates in this species (5).

To accommodate for this grouping behavior, we extended the Vicsek model to include aggregation dynamics as Time-Inhomogeneous Markov Chain process, with probability of aggregation and probability of leaving the aggregate to be dependent on the number of simulated isopods (agents) within a specific interaction radius and by their distance from the edge. In practice, the probability of aggregation was steeply increasing for the number of neighboring agents to change from 0 to roughly 8, and it also showed steep increase within a ring of about 1/10 of arena radius. The probability of leaving the aggregate was smoother, with only dependence of arena radius, dropping by about 50% when moving from the center of the arena to its edge, S1 Fig. We did not aim to fit these parameters but rather wanted to get effective description of trends observed in the experimental data, without overcomplicating the model. With these additions the extended Vicsek model predicted the isopods dynamics over long times, and in specific cases (C, C+L, H) also at short times.

Nevertheless, the model currently does not take into account the effect of different stimuli for increasing the speed of isopods upon stimuli activation and the subsequent relaxation of this speed. First, the control condition and condition with only light have the lowest average speed of isopods shortly after stimuli activation across all experimental conditions, Fig. 2. Second, the combined effect of light and vibration increases the speed at the start of the stimuli exposure more than light or vibration alone, thus suggesting that combinations of stimuli represent a more stressful environment for the isopods, which is in accordance with studies in other members of Malacostraca (41–44) and soil arthropod *Folsomia candida* (45); as well as with the prevalence of multiple stressor synergistic effects in food webs (46). Finally, the relaxation of isopod speed to the levels expected for control can take from several seconds to over a minute, depending on condition, indicating that isopods need longer times to adapt to specific combinations of stimuli, Fig. 3. While the modeled aggregation dynamics drive down the average speed of agents over time, the model does not take into account the variations in relaxation speed. One possible way to introduce these effects in the extended model would be to have the speed of agents v_0_ to be stimuli dependent and with possible time-dependence. Another alternative is to sample initial speed of agents from experimentally obtained distributions of speed and investigate the impact of model with varied speeds on the relaxation.

Other possible extensions of both experimental setup and the proposed modelling framework can be considered given previous work on behavioral ecology. For instance, in *A. vulgare* a degree of habituation in their aggregation dynamics was reported (15), that can be included as a stabilization/habituation term in the probability of aggregation. 3D tracking of large swarms of other invertebrates, such as midges, has shown that they have a high capability to withstand external perturbations (47). However, the role of vibrations for the aggregation dynamics is not clear. Many individuals decided to aggregate more often under the combined stimuli, rather than flee. Taking into account that in *A. vulgare*, isopods chose to aggregate where aggregates were already present (15), and the high sensitivity to substrate-borne vibrations reported in *A. officinalis* (3), modeling aggregation based on local group size and directional noise seems appropriate. However, differences in aggregation dynamics and aggregate sizes in *P. scaber* and *P. pruinosus* in combined stimuli setups have also been reported (5), further complicating faithful modeling of their behavior. Another problem to further refine modeling is the lack of information regarding leadership fluctuation in terrestrial isopods. Here, we assume equal interaction *radii* and agent size affecting the average heading update, producing a probability density map of spatial occupancy (Fig. 6) that ignores intraspecific variations in body morphology, which is known to influence desiccation rate in terrestrial isopods (48), which in turn influences aggregation (49,50).

Regarding limitations, to better understand aggregation dynamics, it is necessary to test the same setups with single-isopod, two-isopod and three-isopod variants. This could yield interesting results that would reveal how the first principles of aggregation work in terrestrial isopods. Additionally, a gradient of light and vibration intensities could be tested to further develop a method based on experimental stimuli-modulated speed sampling. Since non-symmetric setups produced significantly larger effects (one quadrant vs half, and three quadrants vs four quadrants), testing other single-quadrants or diagonal setups could also bring interesting results regarding their response to non-symmetric multi-modal stimuli exposure. From the modeling perspective, another point of improvement could be modeling attractor points: the QAM grips that hold the Petri dish in place seem to create sinks where isopod behavior collects. Additionally, this extension of the model assumes a circular interaction radius, but as noted in earlier works on the Vicsek model, not all agents will display such interaction geometry (51); terrestrial isopod anatomy suggests a more complex sensory function which has been simplified to a circular interaction radius. Collective behavior models constructed using radial basis functions with spatiotemporal pigeon flight data show that a distance-dependent attraction/repulsion rule is enough to reproduce their dynamics (52), so a future direction could be to fit the system with the collected terrestrial isopod data, and measure whether the aggregation dynamics modeled in this work agree with such model. Moreover, it was shown in prawns, that simple memory-less models with self-propelled agents fail to capture the fine scale dynamics of the system, suggesting that models with memory reproduce both the fine-scale and global dynamics much better (53). Another extension of the model could consider social interactions, that were indicated as key factor for driving the collective energetic efficiency in ants (54). Finally, this experiment did not take into account RH%, temperature, or sex-specific behaviors, which are all factors that affect terrestrial isopod behavior (14,55–58).

Nonetheless, this extended model provides a controlled and interpretable framework for testing multi-modal stimuli on terrestrial isopod swarm behavior. It captures behavioral interactions, offering insights for bio-inspired robotics and adaptive collective systems. By comparing model outputs under varying parameters against newly obtained experimental data, we can understand better their responses to light and vibration stimuli. Terrestrial isopods are valuable members of the ecosystem due to their participation in nutrient cycling and their ability to act as biomarkers (59). Another member of the *Porcellionides* family, *Porcellionides cingendus*, has shown potential for environmental assessments (60). This highlights that understanding their response to different stimuli could also provide a better insight into ecosystems engineering.

## Acknowledgements

This research was supported by the strategic programme Excellence Initiative at Jagiellonian University, Poland (RSM/122/FP). PVF and MZ were supported by a grant from the Priority Research Area DigiWorld under the Strategic Program Excellence Initiative at Jagiellonian University and by National Science Center Poland (2021/42/E/NZ2/00188).

## Supporting Information

### Tables

**S1 Table.** Experimental conditions showing the stimuli setup for the 3^rd^ minute. 10 repetitions were done with a swarm size of 25 individuals. C individuals stayed 3 minutes within the enclosure with no stimuli activation. For all other conditions stimuli was activated after 2 minutes and lasted for 1 minute. The activation of stimuli corresponds to *t* = 0 s for the analyzed videos. C+L individuals had the light turned on. H and H+L had half of the actuators turned on. Q and Q+L had only one vibrating quadrant active. 3Q and 3Q+L had ¾ of the actuators working. F and F+L had the full set of quadrants active.

| Condition | n | Light | Q1 | Q2 | Q3 | Q4 | Pictogram |
| --- | --- | --- | --- | --- | --- | --- | --- |
| C | 25 | OFF | OFF | OFF | OFF | OFF |  |
| C+L | 25 | ON | OFF | OFF | OFF | OFF |  |
| H | 25 | OFF | OFF | OFF | ON | ON |  |
| H+L | 25 | ON | OFF | OFF | ON | ON |  |
| Q | 25 | OFF | OFF | OFF | OFF | ON |  |
| Q+L | 25 | ON | OFF | OFF | OFF | ON |  |
| 3Q | 25 | OFF | OFF | ON | ON | ON |  |
| 3Q+L | 25 | ON | OFF | ON | ON | ON |  |
| F | 25 | OFF | ON | ON | ON | ON |  |
| F+L | 25 | ON | ON | ON | ON | ON |  |

**S2 Table.** List of parameters used in the simulations. The list of parameters used in both the radially restricted Vicsek model (null model; eq. 1 in the Materials and Methods) and the terrestrial isopod Vicsek model with aggregation dynamics (extended model; eq. 3 in the Materials and Methods).

| Parameter | Null model | Extended Model |
| --- | --- | --- |
| $v_0$ | 1.0 cm/s | 1.0 cm/s |
| $\eta$ | $0.3 \pi \approx 0.9425$ | 0.9425 |
| $R_{\text{inter}}$ | 1.0 cm | 1.0 cm |
| $p_A^{\text{max}}$ | - | 0.75 |
| $p_L^{\text{max}}$ | - | 0.25 |
| $\lambda_{\text{edge}}$ | - | 3.0 |
| $g_A$ | - | 4.2 |
| $\alpha_{\text{edge}}$ | - | 20.0 |
| $\beta_{\text{edge}}$ | - | 0.92 |

**S3 Table.** The statistics of discarded data points for the relaxation time and probability distribution analysis. Speeds over 8 cm/s were regarded as motion tracking artifacts and discarded from the analysis in Fig. 2, 3, 4 and 5. The columns indicate, from left to right: experimental conditions (see Fig. 1D or S1 Table), total number of speeds collected by the tracking algorithm, number of discarded speeds, relative number of discarded speeds. We additionally note that the maximal possible amount of tracked data points is roughly 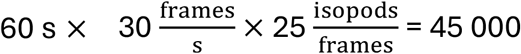 isopods positions in 1 minute, however as isopods localize in aggregates the number of tracked points goes down.

| Condition | Discarded | Total | (%) |
| --- | --- | --- | --- |
| C | 270 | 23809 | 1.13 |
| C+L | 188 | 19771 | 0.95 |
| H | 186 | 15397 | 1.21 |
| H+L | 289 | 19219 | 1.50 |
| Q | 282 | 14524 | 1.94 |
| Q+L | 312 | 13586 | 2.30 |
| Q3 | 216 | 12685 | 1.70 |
| Q3+L | 276 | 12431 | 2.22 |
| F | 193 | 14335 | 1.35 |
| F+L | 267 | 12484 | 2.14 |

### Figures

**S1 Fig.**
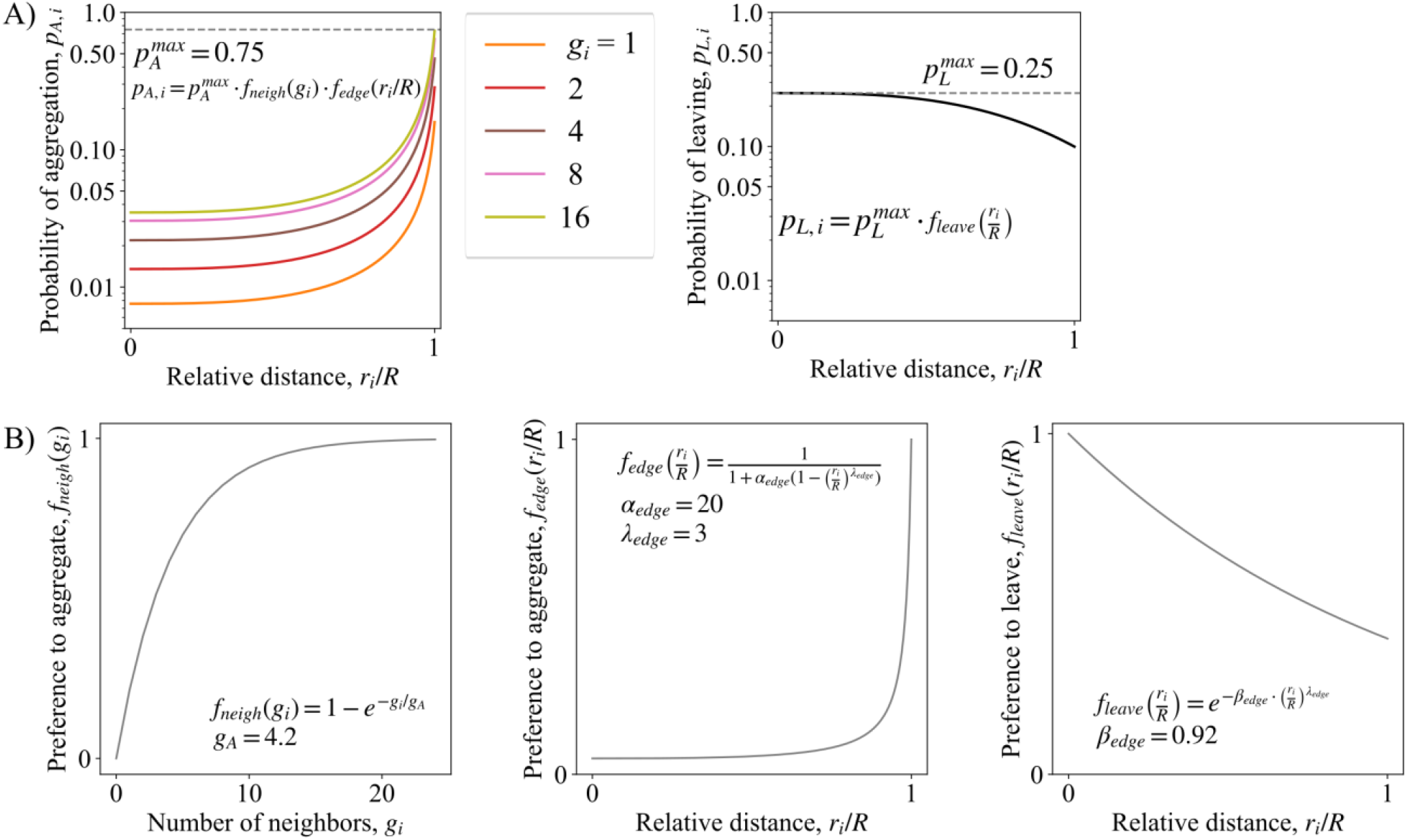
Functions affecting aggregation probability. **A)** The probability of aggregation *p*_A,*i*_(*t*) shows its dependence on the preference to aggregate *f*_*neigh*_(*g*_*i*_) as a function of number of agents *g*_*i*_ within interaction radius and *f*_*edge*_(*r*_*i*_/*R*) function that increases the probability of agents to aggregate near the edge of the circular arena of radius *R*. The probability of leaving *p*_L,*i*_(*t*) depends on the preference to leave the aggregate, *f*_*leave*_(*r*_*i*_/*R*). **B)** The three functions, *f*_*neigh*_(*g*_*i*_), *f*_*edge*_(*r*_*i*_/*R*) and *f*_*edge*_(*r*_*i*_/*R*), that *p*_A,*i*_(*t*) and *p*_L,*i*_(*t*) depend on, with the parameters used for the simulations. See text following eq. 6 and eq. 7 in Materials and Methods for detailed explanation.

**S2 Fig.**
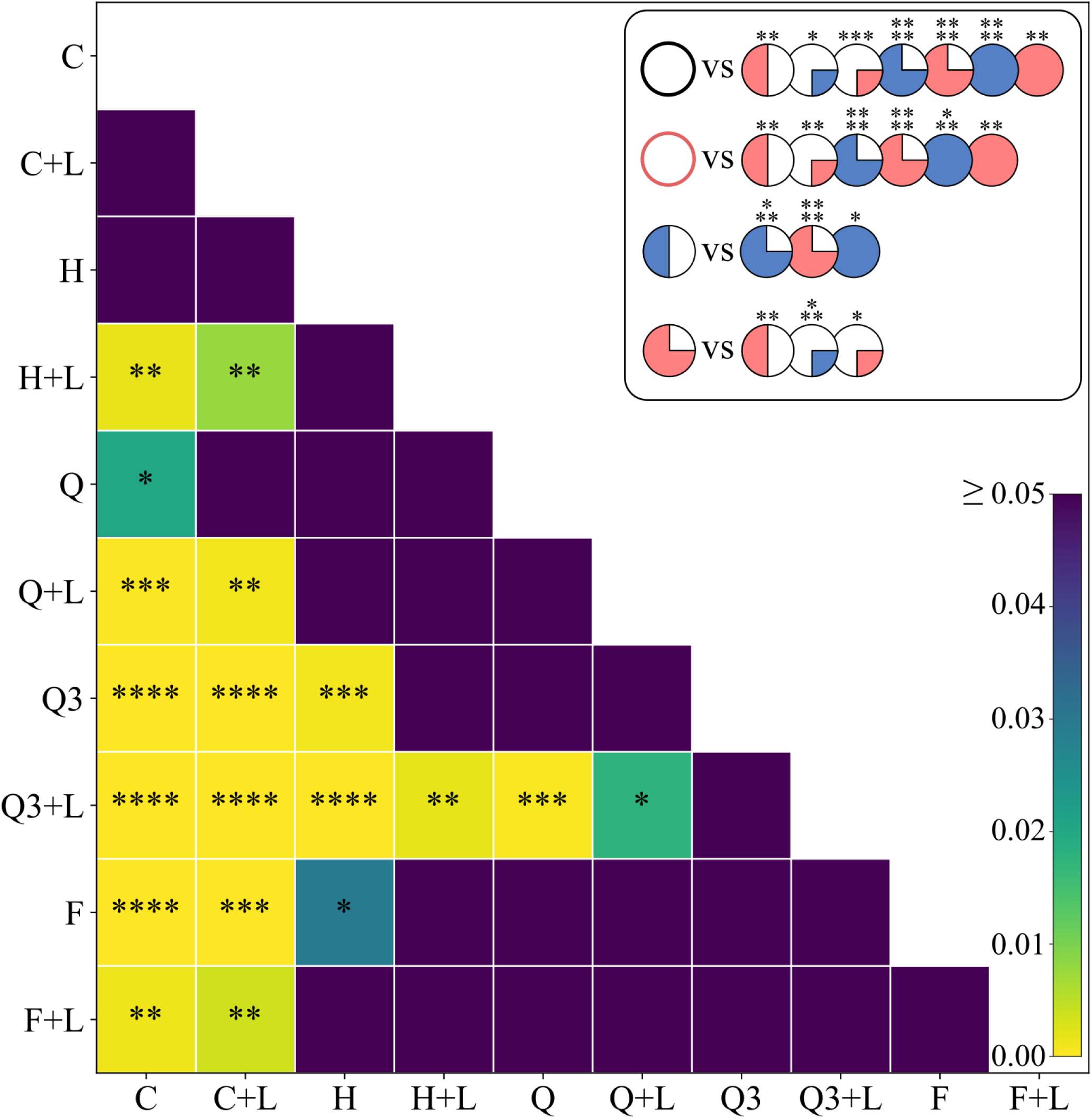
Comparison of statistical significance between early isopod speeds for all conditions. Pairwise Welch independent t-test of tracked speed of isopods at *t* = 2 s after stimuli activation for all conditions. The test statistics for each condition was based on speed estimates that were pooled together for all repetitions for that condition. The color bar represents the holm-adjusted p-value. Significance stars: **** p < 0.0001; *** p < 0.001; ** p < 0.01; * p < 0.05. Non-significant values (p ≥ 0.05) are not shown. The inset shows the significant differences using the pictograms for each setup. See Fig. 1D for an explanation of the pictograms.

**S3 Fig.**
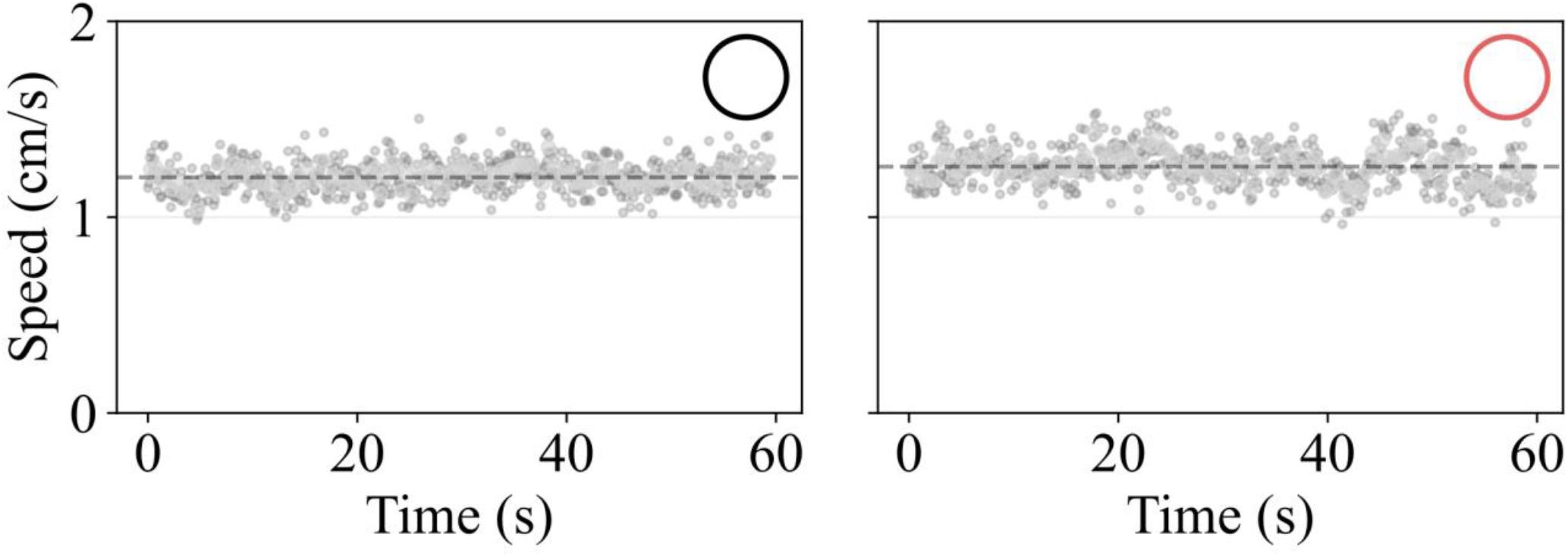
Speed across individuals across repetitions for control (C) and control with light (C+L) conditions. The mean speed of isopods sampled in 0.5 s bins (120 bins in 60 s, 15 frames per bin; gray points) after stimuli activation is indicated. The overall mean speed averaged over 60 s for control condition is 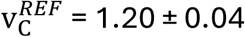 (mean ± SEM; dashed line), and 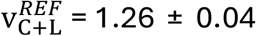 (mean ± SEM; dashed line) for control with light is shown. Note that compared with Fig. 4 there is no relaxation of speed of isopods.

**S4 Fig.**
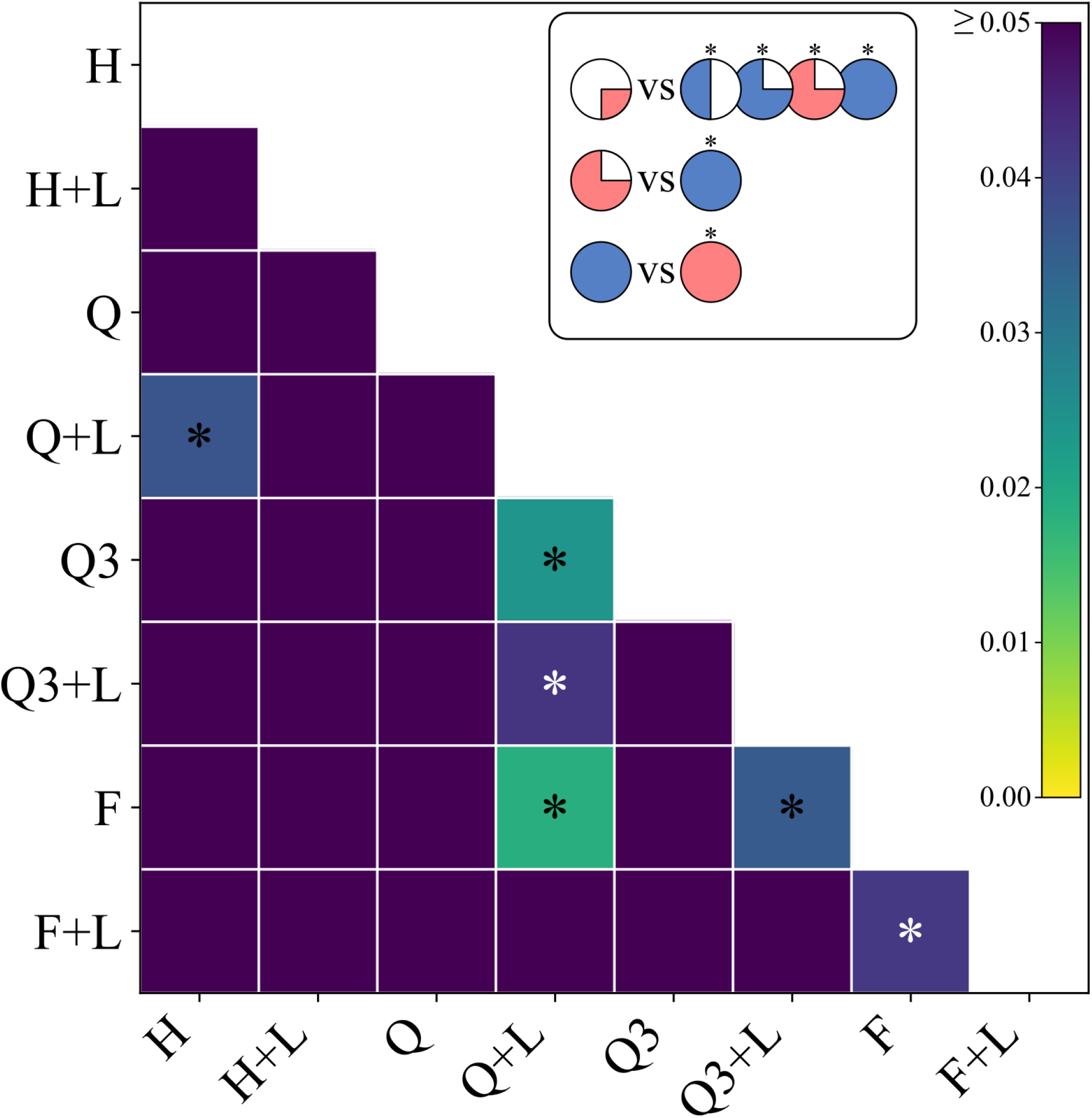
Comparison of statistical significance of log of speed of relaxation time for all tested conditions. For each experimental repetition the mean speed of isopods over 60 s time window, sampled in 0.5 s bins after stimuli activation was collected. An exponential curve of the form v(*t*) = v_∞_ + (v_0_ − v_∞_)*e*^−*t*/τ^ was fit separately for each repetition and condition, where v_0_ was set to the mean speed of isopods in the first bin (*t* from 0 to 0.5 s), and v_∞_ and τ were estimated from the fit. The test statistics for each condition was based on log(τ) estimates for each of the 10 repetitions. The p-value Pairwise Welch independent t-test between any two conditions is indicated by color bar. Significance stars: **** p < 0.0001; *** p < 0.001; ** p < 0.01; * p < 0.05. Non-significant values (p ≥ 0.05) are not shown. See Fig. 1D for an explanation of the pictograms.

**S5 Fig.**
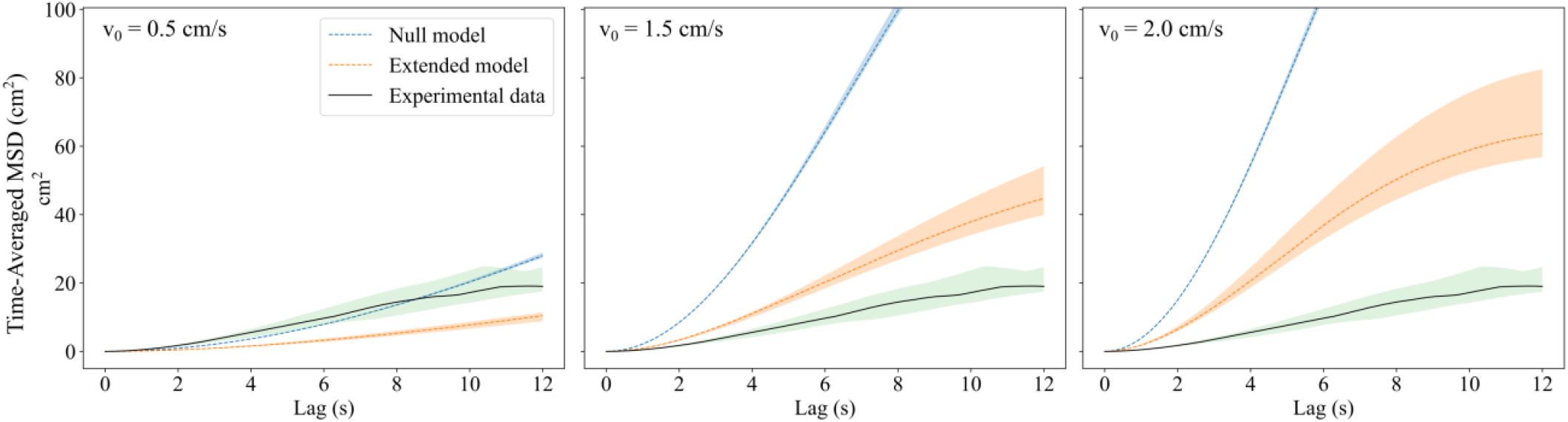
The Time-Averaged Mean Square Displacement for models with different speed of agents against experimental data for control condition. Halving the speed of agents v_0_ allows the null model (blue) to match the experimental data (black, with green confidence bands) better than for the default v_0_ = 1 cm/s, compare with Fig. 7. The extended model (orange) with halved v_0_ underestimates experimentally obtained 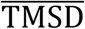. Increasing the v_0_ results in a steep increase of 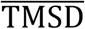 for the null model. The extended model also exhibits a steep increase with increasing v_0_, yet it shows saturation of displacement that results from aggregation dynamics. The bands represent the upper and lower bound quantiles (0.75 and 0.25, respectively).

**S6 Fig.**
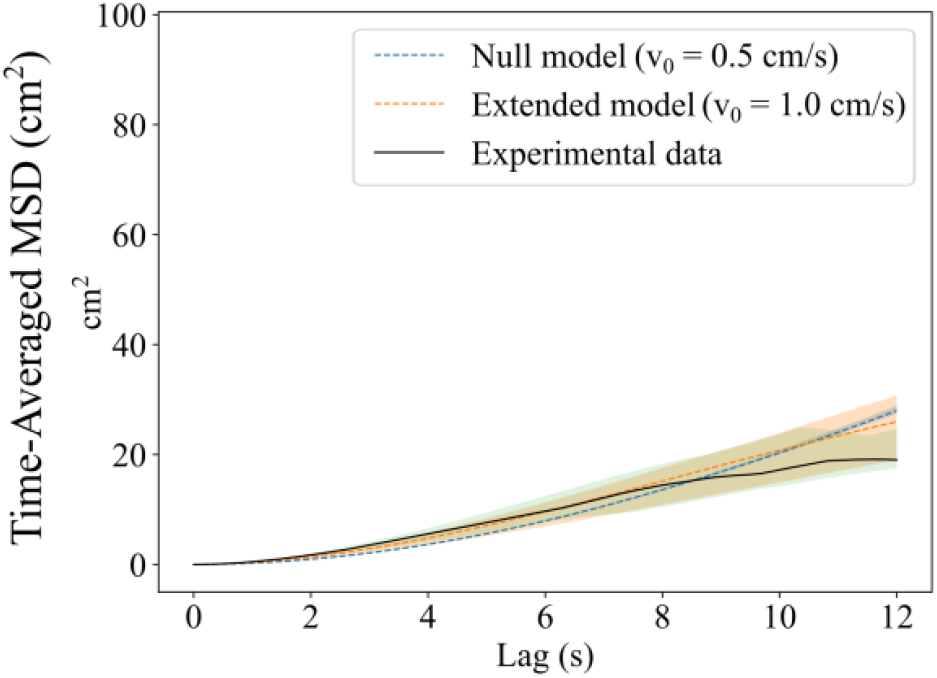
The Time-Averaged Mean Square Displacement for the null model and the extended model with speed of isopods adjusted to match control condition. Both models match the experimental data for the control condition with their respective v_0_ values, see Eq. 1 and Eq. 3. In the case of the null model (blue), halving the speed to v_0_ = 0.5 cm/s allows the model to match the experimental data (black, with green confidence band) better, than the default v_0_ = 1 cm/s, see Fig. 7. However, the null model spatial occupancy with v_0_ = 0.5 cm/s is far from experimental counterpart, compare S7 Fig with Fig. 5. The extended model (orange) needs a constant speed of 1.0 cm/s to match the experimental control and qualitatively reproduces spatial occupancy. The bands represent the upper and lower bound quantiles (0.75 and 0.25, respectively).

**S7 Fig.**
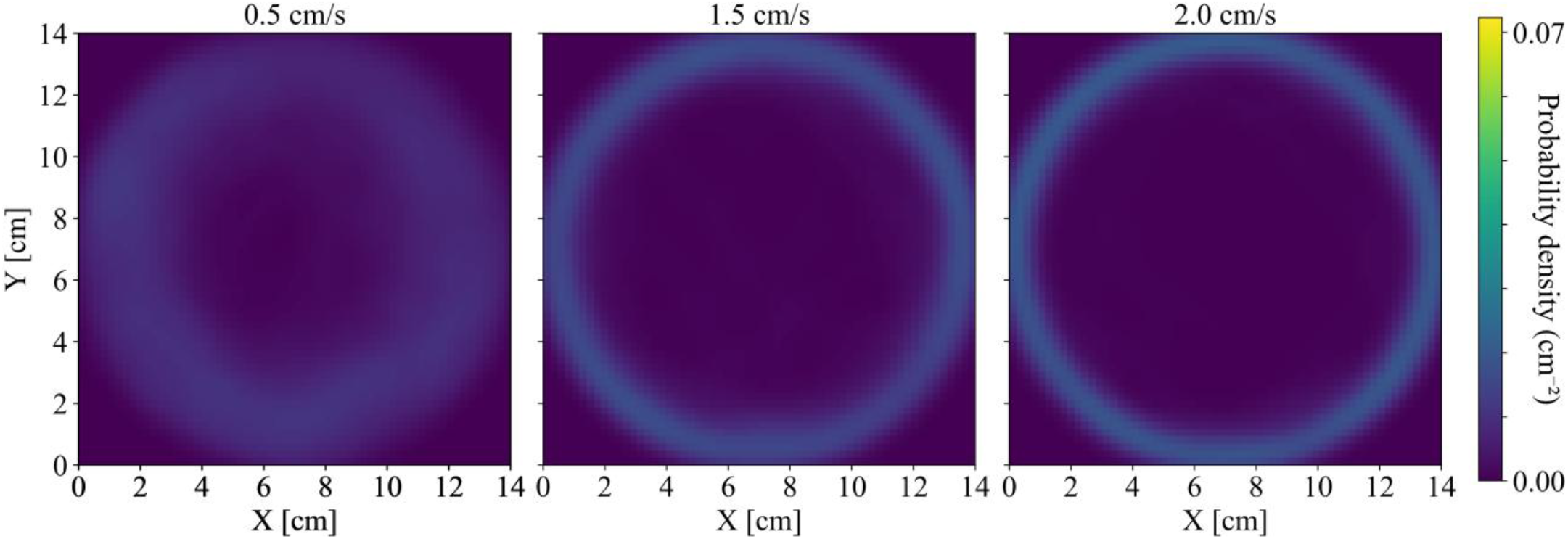
Probability density of spatial occupancy of agents in the null model with different agent speeds. Left: null model data KDE with v_0_ = 0.5 cm/s. Center: null model data KDE with v_0_ = 1.5 cm/s. Right: null model data KDE with v_0_ = 2.0 cm/s. See Fig. 5 for the spatial occupancy of the experimental data.

**S8 Fig.**
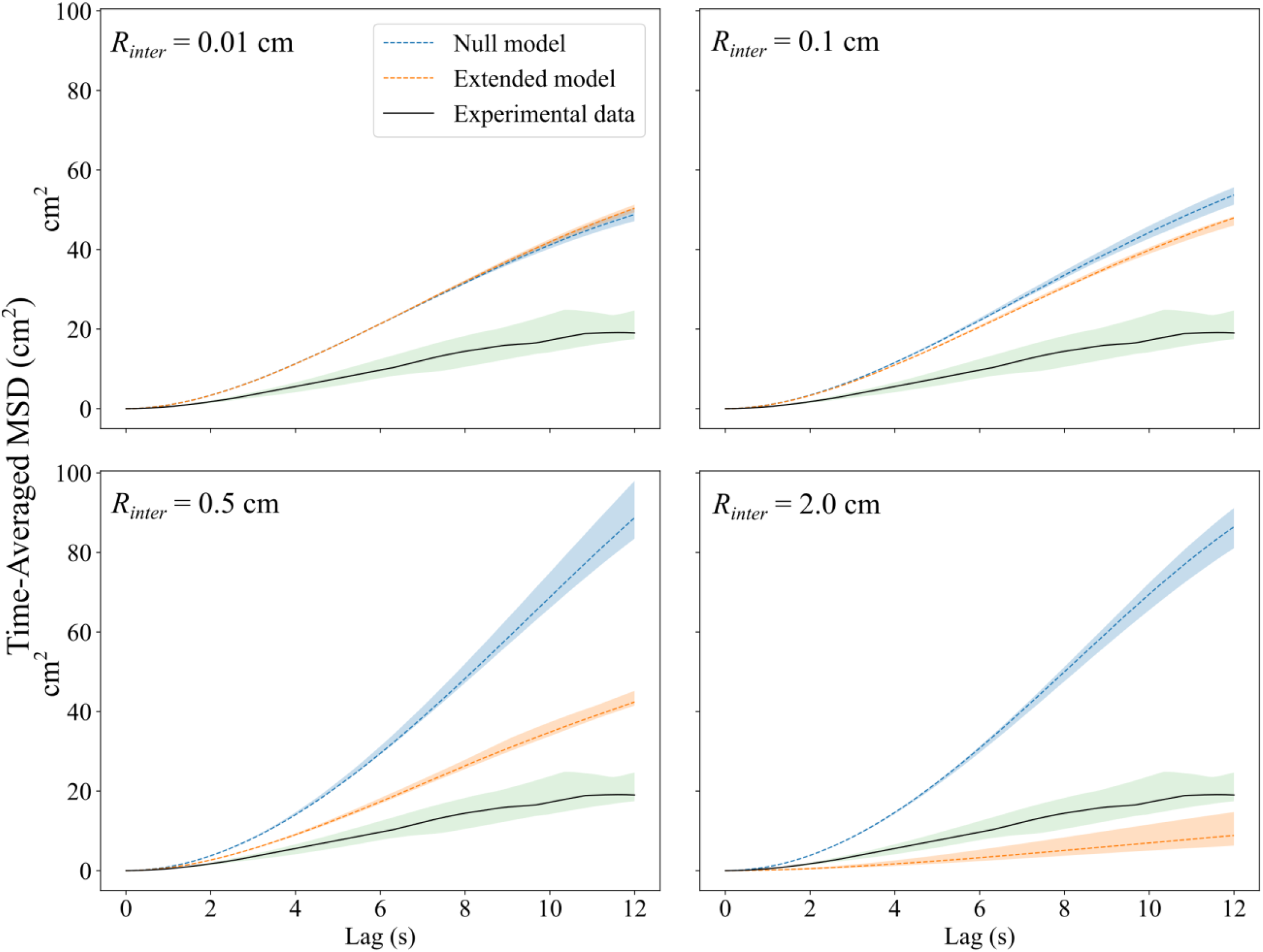
Time-Averaged MSD for null models with different radii values against experimental data (Control) and the extended model. Even after reducing the radius of interaction of agents to values that are biologically unreasonable for terrestrial isopod interaction, the null model (blue) does not approximate the isopods’ 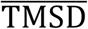 measurements (black, with green confidence bands). As *R*_*inter*_ increases, agents are more likely to aggregate. At *R*_*inter*_ = 2 cm the extended model (orange) drives the 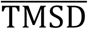 down, which is consistent with an increased aggregation tendency, resulting from increasing number of neighbors. The bands represent the upper and lower bound quantiles (0.75 and 0.25, respectively).

